# Fishing for a reelGene: evaluating gene models with evolution and machine learning

**DOI:** 10.1101/2023.09.19.558246

**Authors:** Aimee J Schulz, Jingjing Zhai, Taylor AuBuchon-Elder, Mohamed El-Walid, Taylor H Ferebee, Elizabeth H Gilmore, Matthew B Hufford, Lynn C Johnson, Elizabeth A Kellogg, Thuy La, Evan Long, Zachary R Miller, M Cinta Romay, Arun S. Seetharam, Michelle C Stitzer, Travis Wrightsman, Edward S Buckler, Brandon Monier, Sheng-Kai Hsu

**Affiliations:** Section of Plant Breeding and Genetics, Cornell University, Ithaca, NY USA 14853; Institute for Genomic Diversity, Cornell University, Ithaca, NY USA 14853; USDA-ARS; Ithaca, NY, USA 14853; Department of Computational Biology, Cornell University, Ithaca NY USA 14853; Donald Danforth Plant Science Center, St. Louis, MO USA 63132; Department of Ecology, Evolution, and Organismal Biology, Iowa State University, Ames, IA USA, 50011

**Author notes:** co-corresponding authors; Corresponding authors: Aimee J Schulz and Brandon Monier. co-first authors (Aimee J Schulz, Sheng-Kai Hsu).

## Abstract

Assembled genomes and their associated annotations have transformed our study of gene function. However, each new assembly generates new gene models. Inconsistencies between annotations likely arise from biological and technical causes, including pseudogene misclassification, transposon activity, and intron retention from sequencing of unspliced transcripts. To evaluate gene model predictions, we developed reelGene, a pipeline of machine learning models focused on (1) transcription boundaries, (2) mRNA integrity, and (3) protein structure. The first two models leverage sequence characteristics and evolutionary conservation across related taxa to learn the grammar of conserved transcription boundaries and mRNA sequences, while the third uses conserved evolutionary grammar of protein sequences to predict whether a gene can produce a protein. Evaluating 1.8 million gene models in maize, reelGene found that 28% were incorrectly annotated or nonfunctional. By leveraging a large cohort of related species and through learning the conserved grammar of proteins, reelGene provides a tool for both evaluating gene model accuracy and genome biology.

## Introduction

The number of available plant reference genomes has reached close to 800 species over the past 20 years, with a 50-fold increase in the last decade alone^1,2^. In the near future, we will likely see sequencing of the majority of plant species^3^. The increasing availability of genome assemblies has enabled significant advances in genetics and genomics, especially in non-model species that previously had limited resources.

Despite this tremendous increase in genome assemblies, methods for gene model annotation have not scaled with these advances^4^. Gene model annotation, while seemingly straightforward, is quite challenging in practice as it involves the modeling of the entire central dogma of molecular biology, and current methods tend to over-annotate genes. *Ab initio* gene model prediction methods are trained to look for specific signals and motifs that are known to be characteristic of a protein coding gene (i.e, splice sites, coding sequences, and transcription/translation start/stop sites). Other methods of gene annotation include evidence-based predictions of transcriptomes from datasets including RNAseq, cDNA, expressed sequence tags (ESTs), and diverse protein databases. In a recent benchmarking study^5^, AUGUSTUS^6^, which combines *ab initio* and evidence-based methods, was the most accurate method but still had 76.5% of all predicted proteins incorrectly predicted. Gene prediction errors propagate down the central dogma, as found by Bányai et al^7^, where previous reports of proteome innovation in lancelets were shown to be false, and instead were attributed to gene prediction errors. While users of genome browsers can flag specific gene models with annotation errors (266 genes have had errors identified by users on MaizeGDB^8^) and additional manual curation efforts have been published (i.e., in one study over 900 undergraduates curated Muller elements^9^), this approach of manual curation is both uncommon and time intensive.

Errors and variation in gene models can stem from both technical and biological sources. Technical sources can result from using models trained on other species^10^ and from misalignment of reads, causing induced insertions, deletions, or gene splitting^11–13^. The nature of sequencing mRNA itself for evidence-based gene model annotation can result in errors, simply from the sequencing of pre-spliced mRNA transcripts^4,14^. Gene annotation is further complicated by sources of biological error, such as transposable elements. In particular, Helitron elements pick up and disperse gene fragments^15^, generating tens of thousands of transcribed sequences^16^. Additionally, transposon-encoded genes that produce proteins necessary for transposition are often incompletely masked by gene annotation pipelines, and transposable element-encoded genes can then be included in annotated genes^17,18^.

To sort through possible annotation errors and reduce the number of over-annotated genes, we propose leveraging the power of evolutionary constraint to identify the most conserved and functional gene models using machine learning. The strongest degree of evolutionary constraint can be observed at the level of the proteome (Figure 1A). This derives from the cellular energetic cost of translation, where over 99% of the energy spent goes towards translating the protein, while 0.1% goes towards transcribing the gene into mRNA, and 0.03% towards replicating DNA^19^. The evolutionary constraint on translated proteins has created a learnable grammar for their structure and folds that is consistent across all of life^20,21^. Though under less constraint, we also expect a certain learnable grammar for mRNA transcripts (e.g., their coding sequence, splicing sites) among closely related species. We have developed an approach, reelGene, for the **<u>R</u>**obust **<u>E</u>**valuation of gene model annotations using **<u>E</u>**volutionary signals **<u>L</u>**everaged at both transcriptional and translational levels. As a case study, we have evaluated over 1.8 million gene models across 26 diverse maize assemblies^22^. Maize provides an excellent study system due to the complexity of its genome and resulting large number of gene models, relatively high levels of genetic diversity, and gene presence/absence variation identified across the 26 diverse lines. Our approach reels in the most evolutionarily supported genes from a given annotation, generating a more robust working set of genes, and greatly helps future genetic studies by removing genes less likely to make a functional protein.

**Figure 1.**
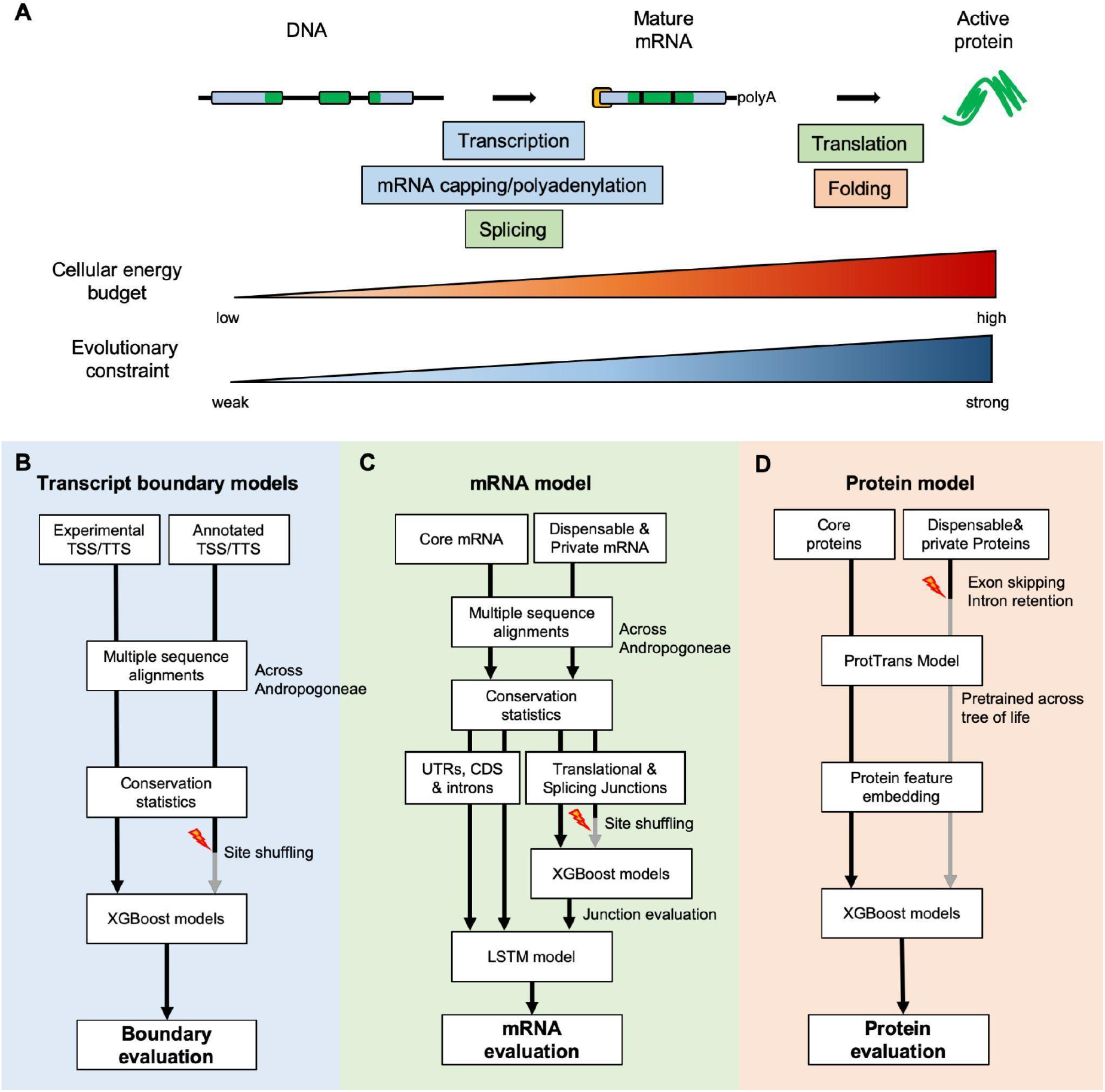
Graphic summary of reelGene models. A. The energy cost and evolutionary constraint increases across the central dogma. Given the variation, we developed three submodels to evaluate different components of the process: **B.** Transcript boundary models were trained to learn the conservation pattern of experimentally validated transcription start and termination sites (TSS and TTS); **C.** The mRNA model integrates the conservation of multiple components of a mRNA transcript and is trained to recognize the sequential grammar of a core gene model shared by 26 diverse maize lines; **D.** The protein model builds on feature embedding from an unsupervised machine learning model pre-trained across the tree of life and learns the folding principles of the core proteins conserved across 26 diverse maize lines.

## Results

The reelGene framework is built upon three focus areas – conservation of transcript boundaries, mRNA sequence, and protein sequence – resulting in three different submodels (Figure 1B-D). Since the local interactions between amino acids within a peptide are constrained by their physicochemical properties, we are able to leverage a protein language model pre-trained on prokaryotic and eukaryotic data^20^ to evaluate evolutionarily supported proteins. In contrast, mRNA transcripts are under less constraint, which requires data from closely related species. In the case of maize, we looked to the encompassing Andropogoneae tribe.

Andropogoneae includes roughly 1,200 species of C4 grasses that shared a common ancestor 16-20 million years ago and came to dominate most tropical and some temperate grasslands ^23,24^. We leveraged 33 *de novo* long read assemblies^8^, and sequenced and assembled the genomes of 58 additional Andropogoneae species using short reads, totaling 91 species. We were able to assemble most genes at full length, a subset of those including multiple copies, but some genes remained fragmented and split across multiple contigs. TABASCO^25^ scores (a grass-specific BUSCO-like evaluation for 5,600 conserved genes) for the short read assemblies averaged at 70% complete, 16% complete and duplicated, 13% fragmented, and less than 1% missing. We used these species to generate multiple sequence alignments to each maize transcript, totaling 1.8 million transcripts across 26 maize lines^22^. On average, each MSA consisted of alignments to 61 species (supplementary fig S1), capturing close to 500 million years of evolution across the panel (supplementary fig S2), and allowing on average 4 independent mutations at each neutral site.

Transcriptional initiation and termination determine the regulation and stability of an mRNA transcript, and these transcriptional boundaries make up the first model of reelGene. Incorporating experimental evidence ^26,27^ as positive sets, we trained two separate XGBoost^28^ classifiers to identify transcriptional start and poly-adenylation sites in our multiple sequence alignments (Figure 1B). We simulated false transcript boundaries by base-pair shuffling empirical boundaries during model training (negative sets). Evaluating on 5,000 random left out genes, these models perform moderately well (Start Model: ROC-AUC of 0.848 and PR-AUC of 0.325; Stop Model: ROC-AUC of 0.878 and PR-AUC of 0.547; Supplementary figure S2). Potentially, the moderate performance reflects (1) the limitation on the available training datasets and/or (2) the weaker selection constraint on transcript boundaries^26^.

Our second model encompasses a two-layer machine learning model that investigates the sequence conservation of mRNA transcript elements from multiple sequence alignments (Figure 1C). The mRNA model includes four XGBoost classifiers for translational start, translational stop, splicing donor and splicing acceptor junctions, and an LSTM model that incorporate the conservation of 5’ UTRs, coding exons, introns, and 3’ UTRs. As with our first model, we simulated false junctions by the base-pair shuffling approach, enabling effective recognition of the conserved motifs and grammar across different elements (supplementary figures S3 and S4). Using a leave-one-chromosome-out (LOCO) approach, we trained our mRNA model on transcripts shared across 26 maize lines (core; positive set) and those shared across 1-23 of the lines (dispensable or private; negative set). We assumed that genes not shared across all 26 maize lines were less likely to be functionally conserved proteins. The mRNA-based approach achieves high precision and accuracy in distinguishing between maize core and noncore genes, with an average ROC-AUC of 0.927 and PR-AUC of 0.910 across all 10 models (supplementary figure S5). Testing the models on 5,000 withheld maize genes, the AUC reaches 0.946 and 0.989 for ROC and PR curves, respectively (Figure 2A). These high values highlight the strength of using evolutionary conservation of closely related species to identify functionally conserved gene models.

**Figure 2.**
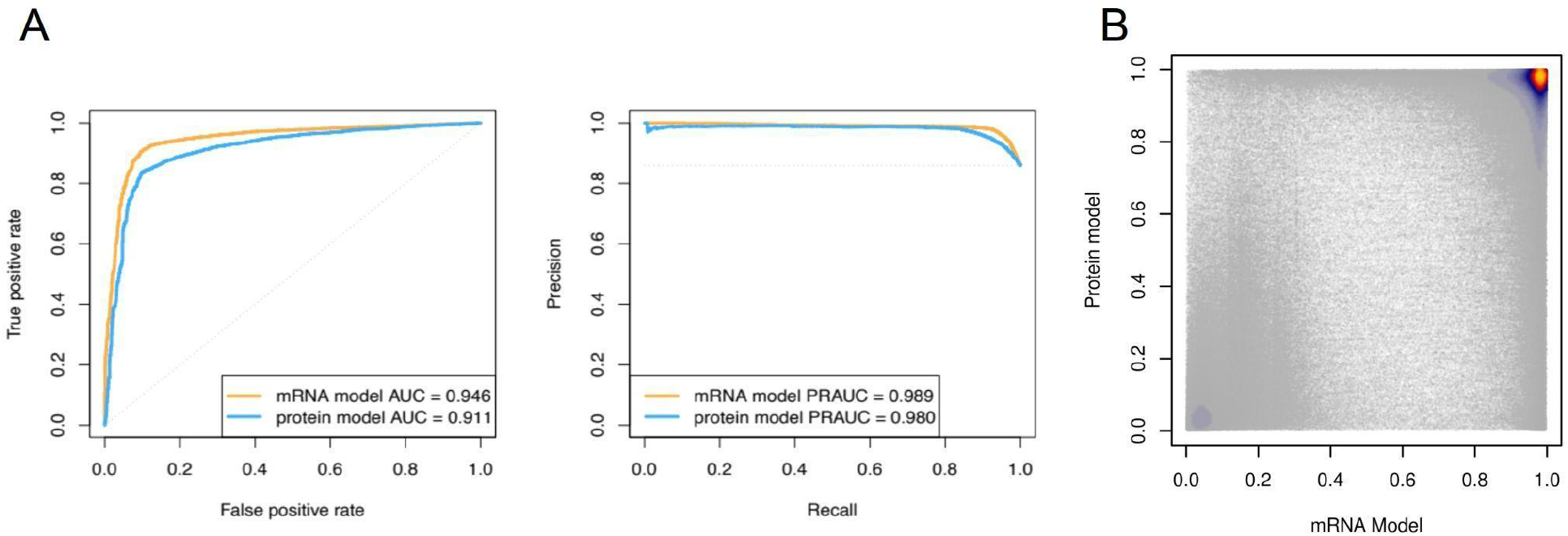
mRNA and Protein model performance. **A**. ROC and PR curves in the mRNA submodel (orange) and the protein submodel (blue) for our 5,000 withheld genes **B**. Scatter plot of the scores from the mRNA submodel (x-axis) and the protein submodel (y-axis). Heat color denotes the density of dots.

In our third model, we utilize the extensive gains in protein prediction of transformer language models^20,21^ to evaluate the evolutionarily conserved grammar found across proteins (Figure 1D). We extracted the embeddings of the 1.8 million maize transcripts from the protein language model ProtTrans^20^, and used them to train an XGBoost model to identify conserved proteins. Like the mRNA model, we trained the protein model on the core and noncore gene sets. Additionally, we simulated both intron retention and exon exclusion as part of our negative set. This addition makes our model more robust and ensures that it learns the correct signatures of a functional protein instead of memorizing the patterns of a core versus noncore gene. Our protein model achieves high levels of precision and accuracy, with an average ROC-AUC of 0.944 and PR-AUC of 0.881 across all 10 models (supplementary figure S5). For the 5,000 withheld maize genes, the ROC-AUC is 0.911 and the PR-AUC is 0.980 (Figure 2A). The error rate of classification was observed to be independent of the number of exons in a gene. Because this model does not rely on a direct comparison of sequence to other species, it is suitable for investigating genes that are only found in a few species.

The protein and mRNA-based models agree moderately well (Pearson = 0.707, Spearman = 0.511, Figure 2B). The mRNA-based model can detect all essential elements of a transcript. For example, some transcripts that do not start with an “ATG” score poorly with the mRNA-based model (<0.01) and well with the protein model (>0.9) look to be mis-annotated antisense non-coding RNA (e.g. supplementary figure S7). The protein model, however, has the advantage of not relying on alignment to other species, allowing the evaluation of novel genes without bias due to alignment constraints. Given these two distinct strengths, we calculate our reelGene score as the average of these two models (supplementary figure S8). In the case of multiple transcripts, we recommend using the two transcript boundary models to differentiate which transcripts have supported transcription start and stop sites.

Using a threshold of 0.5, reelGene classifies 80.5% of all transcripts and 72% of genes as functional, implying that 28% of annotated maize genes have errors or are not functional genes. This threshold can be adjusted to meet different levels of sensitivity (i.e., the percentage of core genes classified as functional). As multiple functional transcripts are identified in 46% of functional genes, we are likely capturing real cases of alternative splicing. To validate our model, we compared reelGene classifications to two held out datasets – one of carefully curated genes with experimentally-verified gene models and another of a proteomic atlas.

In the maize classical gene set^29^ (genes with at least three publications) we find 99.2% (412/415) of classical genes are classified as functional in the current v5 genome annotation version. The three classical genes with low reelGene scores are *hm2* and two copies of *afd1*. In the case of *hm2*, reelGene appears correct in calling the v5 gene model nonfunctional, as the gene model is truncated compared to previous genome annotations and the cloned gene^30^. The reelGene scores of the two previous genome annotations are classified as functional. For *afd1*, the v3 annotation is nearly 32kb in length, which is spanned by three gene models in the v5 annotation. Two of the three v5 genes are classified as nonfunctional, while the third is barely above the 0.5 threshold. In comparison, the v3 gene model scores as functional (0.91). While *afd1* has been described extensively^31–36^, it has not been cloned. This illustrates that caution is needed when evaluating genes in large, complex areas.

In the case of the proteome, we expect that every gene with proteomic evidence should have at least one functional model. reelGene classified 92.2% of genes that are found in the proteome^37^ (maximum FPKM > 1) as functional in the v5 annotation (Figure 3A), and 87.9% with the v3 gene models used in data collection. The 7.8% of v5 genes classified as non-functional have a mean FPKM of 85.5 versus the average of 160 for the functional genes, suggesting that they could be the result of peptide assignment errors.

**Figure 3.**
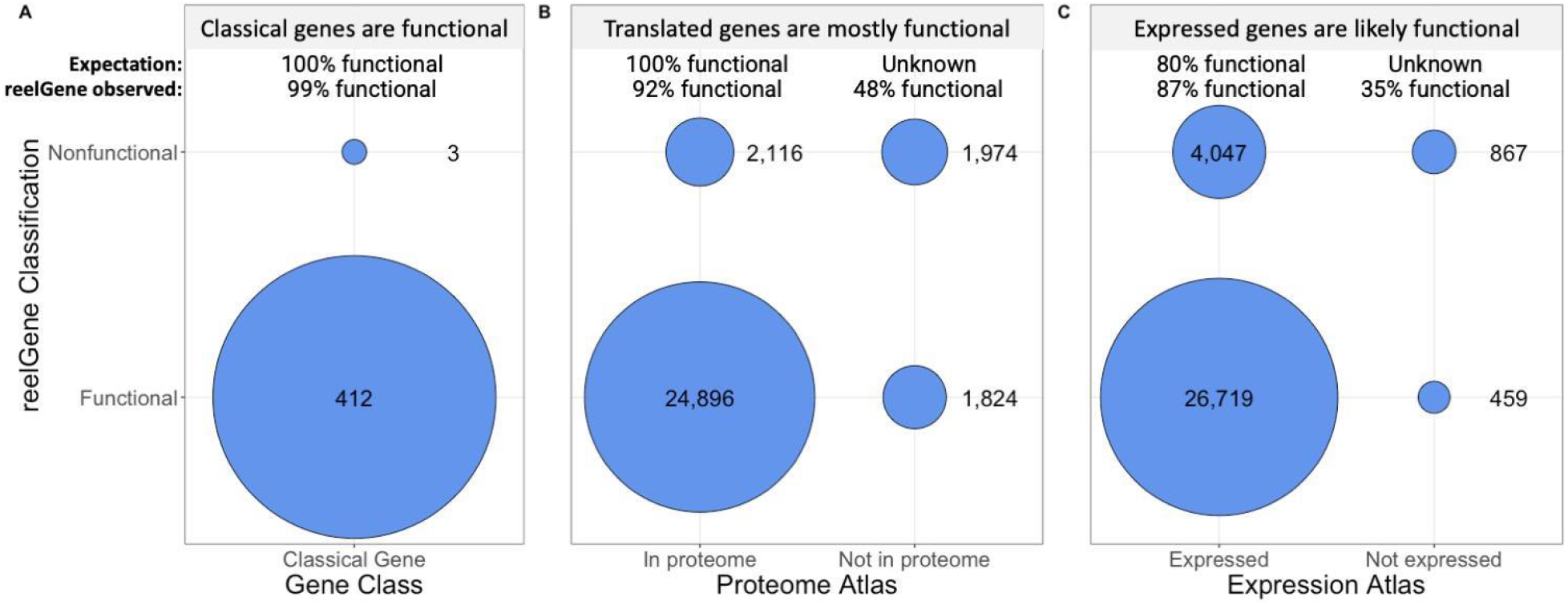
Classification of maize gene models by reelGene. Circles represent the number of genes in each dataset that fall into each category. **A.** reelGene classifies 99% of the classical genes as functional, where we expect 100% to be functional. The 3 genes classified as nonfunctional appear to be annotation errors. **B.** For genes found in the proteome atlas, we expect 100% of them to be functional. reelGene classifies 92% of them as functional and 48% of genes not in the proteome as functional. **C.** Evaluating genes found in the maize expression atlas, we find that 87% of those above baseline expression levels are classified as functional, which is greater than our expectation of 80%. Of the genes that are not in the expression atlas, 35% are predicted as functional.

The correlation between transcription and translation across genes is modest in many species, suggesting that expressed transcripts may not always be translated to functional proteins^38^. Equally likely, difference in molecular turnover rates, cell-type specificity and extraction efficiency could contribute to the discrepancy. Comparing reelGene scores to genes in the maize expression atlas^39^, we find that 86.8% of expressed genes (maximum TPM >1) are classified as functional (Figure 3B), which is above random expectation (80%; the percentage of transcripts classified as functional by reelGene). However, the lower percentage confirms that while correlated, transcriptional activity of a gene does not necessarily translate to protein functionality.

Across maize reference genome (B73) versions, we find that the latest version – v5 – is the most accurate, with 78% (49,043) of all transcripts scoring above the 0.5 threshold, versus 70% (41,921) and 69% (90,157) in v3 and v4, respectively. However, as noted above, occasionally the v5 annotation for a gene was incorrect and the v3 or v4 version scored as functional. reelGene allows users to compare gene models across multiple annotation versions, which will prove invaluable for molecular experiments and gene editing.

In addition to evaluating annotations, reelGene offers insights into genome evolution, population genetics, and molecular biology. In plants, whole genome duplication events occur often and can lead to rapid genome evolution^40^. Maize underwent a whole genome duplication event approximately 12 million years ago, and the genome then fractionated and returned to a diploid state^29,41^. However, thousands of genes are still retained as duplicate copies. reelGene provides new information on the fractionation of the maize genome. In agreement with the maize proteome atlas^37^, genes that are assigned to either of the maize subgenomes are largely classified as functional (97.5%) while less than half (48.9%) of those not assigned are classified as functional. Of the 5,258 gene pairs where copies in both subgenomes are retained, a majority of both copies are functional. In the cases where one copy is functional, and the other is not, we find a significant 30% bias towards the M1 subgenome for retaining functionality (Chi-square test; p-value = 0.011), confirming previous results of preferential retention of M1 genes ^22,29^.

Species or clade specific genes are thought to underlie novel traits which may confer ecological or environmental adaptation ^42,43^. However, identification of such genes has often yielded a long list of candidates and many false positives. Maize and the Andropogoneae have ample opportunity to create novel and clade-specific genes. Of the 1.8 million transcripts evaluated with reelGene, we identified approximately 103,000 genes that have sequences found only within *Zea* (which diverged from *Tripsacum* approximately 0.65MYA^44^), and an additional 15,000 genes that are specific to maize. reelGene suggests that only 2.7% of these 15,000 potentially *de novo* genes are functional (Figure 4). Within B73, reelGene classifies 121 of these young genes as functional. Only 60 of the 121 appear in the maize proteome and are involved in diverse biological processes (supplementary table S1) but whether these genes contribute to local adaptation requires further investigation.

**Figure 4.**
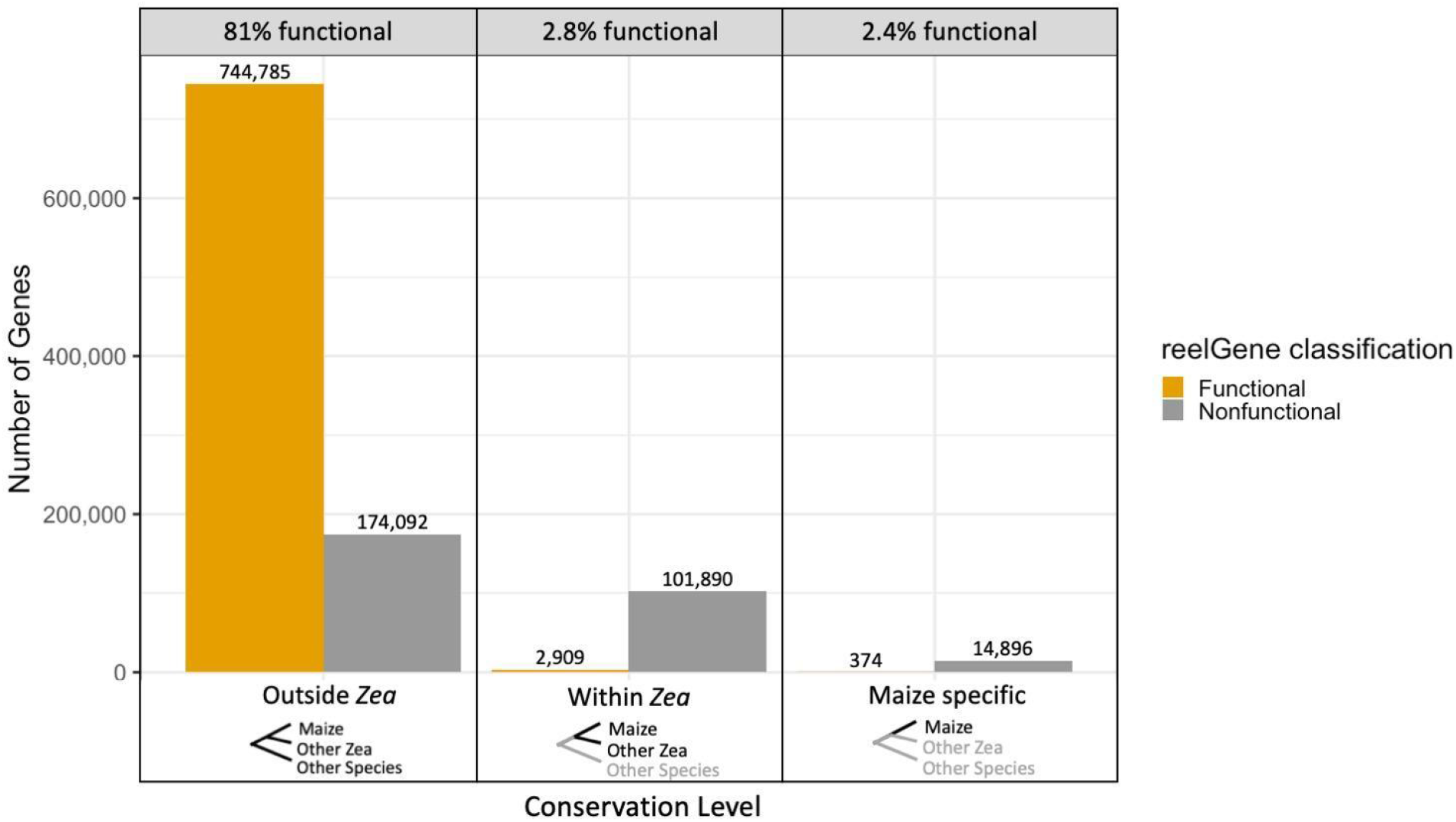
reelGene classification of genes found at varying conservation levels. reelGene classifies most genes that align to species outside of Zea as functional. The vast majority of genes that are found solely within *Zea* or only in maize are nonfunctional.

The maize genome is tremendously diverse (with an order of magnitude more diversity than other species^45^), and the resulting dispensable genes have been hypothesized to be key to heterosis^46,47^ and adaptation^48–50^. Here, reelGene scores 10.3% (793) of the dispensable genes present in B73^22^ as functional. They are enriched in biological processes including defense responses and temperature acclimation (Table 1), reflecting adaptation to local pest/disease and climate challenges. Additionally, recent organelle insertion^51^ appears to contribute to some of these functional dispensable genes (supplementary table S2). Mutations in functional genes could equally contribute to local adaptation and heterosis. In a recent test of multi-location hybrids^52^, protein variants in 281 genes (encompassing both conserved and presence/absence genes) were shown to significantly contribute to heterosis. We find no enrichment of the functional but dispensable B73 genes towards heterosis (Fisher’s exact test; p-value = 0.67). A test of enrichment shows that 96% of heterosis variants are in functional genes and the majority of them are functional across all 26 maize lines which suggests that heterosis can be modulated by conserved functional loci. In a large survey of 5,000 landraces, 186 genes were identified as possibly being involved in abiotic genotype by environment interactions (GxE)^53^. These genes are also enriched for functionality in the reference genome, while the functional but dispensable B73 genes are not, which suggests that, while some biotic GxE may involve segregation of non-functional genes, most abiotic GxE is focused at functional genes.

**Table 1.** Top 10 enriched Gene Ontology terms with fold enrichment > 4 among the dispensable but functional genes.

| GO ID | Term | Annotated | Significant | Expected | p-value | Fold Enrichment |
| --- | --- | --- | --- | --- | --- | --- |
| GO:0009853 | photorespiration | 42 | 5 | 0.64 | 0.0004 | 7.81 |
| GO:0010623 | programmed cell death involved in cell development | 15 | 3 | 0.23 | 0.0014 | 13.04 |
| GO:0017157 | regulation of exocytosis | 18 | 3 | 0.28 | 0.0025 | 10.71 |
| GO:0045492 | xylan biosynthetic process | 70 | 6 | 1.07 | 0.0045 | 5.61 |
| GO:0006904 | vesicle docking involved in exocytosis | 26 | 3 | 0.40 | 0.0072 | 7.50 |
| GO:0006788 | heme oxidation | 9 | 2 | 0.14 | 0.0079 | 14.29 |
| GO:0016102 | diterpenoid biosynthetic process | 54 | 4 | 0.83 | 0.0095 | 4.82 |
| GO:0009631 | cold acclimation | 31 | 3 | 0.48 | 0.0117 | 6.25 |
| GO:2000022 | regulation of jasmonic acid mediated signaling pathway | 58 | 4 | 0.89 | 0.0121 | 4.49 |
| GO:0009306 | protein secretion | 33 | 3 | 0.51 | 0.0139 | 5.88 |

## Discussion

reelGene provides a unique framework to the larger genetics community that allows for the evaluation of gene models through an evolutionary lens. Its potential applications encompass a wide range of research areas. reelGene can help reduce errors in genome annotation by evaluating the conservation of particular key junctions within a gene model. It also enables the evaluation of conservation in genes of interest across any genome, creating opportunities for genome research on the evolution of gene function within a shorter evolutionary time scale. This offers valuable insights into identifying a narrower set of candidate genes with evolutionary support, which is especially relevant to climate change and the need to identify potentially adaptive alleles. Additionally, and perhaps most directly, reelGene predicts whether a gene goes on to make a protein. While it does not replace experimental validation, the strong correlation with protein abundance in addition to near-perfect classification of known functional (Classical) genes points towards reelGene being an appropriate proxy for functionality. Trait-associated variants can be evaluated for functionality, which, in turn, can offer insights into the mode of action (i.e., regulation, protein modification, or sequence gain/loss) underlying specific traits for an organism. reelGene provides another critical link between genotypic and phenotypic variation.

While our pre-trained models immediately lend themselves to many other grass species, the reelGene modeling framework can be adapted to any species of interest. Our protein model is the most straightforward to use for any species, requiring only coding sequences from a genome or transcriptome. The mRNA model is slightly more complicated, but can be replicated by providing genome assemblies of related species and proper training datasets. This will allow a user to assess the functionality of any annotated gene model in their favorite species. However, the biggest challenge when recreating our pipeline with another species is identifying a proper training dataset. One solution, as done in this study, is to use pan-genes identified across multiple accessions of the same species. Another method could utilize pan-genes from multiple, closely related species. Given that we found that species-specific genes are largely non-functional, this provides a robust set of negative training datasets for many species.

As more species are sequenced, the reliance on comparative genomics statistics across species in our mRNA and transcript boundary models could limit the up-scaling of our model, as these steps to generate the multiple sequence alignments and input matrices for our model training are the most time-intensive. This is where the integration of protein language model ProtTrans has significantly enhanced the utility and effectiveness of the reelGene framework and opened up new avenues for protein functionality research. DNA language models are rapidly advancing, but currently, they are mostly pre-trained on human^54^ and vertebrate genomes^55^. Models pre-trained on plant genomes may improve our mRNA and junction boundary models. This shift would empower large language models to autonomously learn the DNA language grammar, presenting an opportunity to expand the scope of applications and enhance the accuracy of our analyses across species.

## Methods

### Sample Collection

Plant material was either a) collected in the field by our team and collaborators, and, b) grown from seed obtained from the Millenium Seed Bank at the Royal Botanic Gardens (RBG), Kew or the USDA-ARS National Plant Germplasm System, or c) obtained as small fragments of herbarium specimens held at the herbarium of the Missouri Botanical Garden (MO), or the herbarium at RBG, Kew (K). Live plant material was maintained in a greenhouse for the duration of the project. Specimens were grown to flowering to verify identity, and voucher specimens were deposited at MO.

### DNA Extraction

DNA was extracted from leaves using a Qiagen kit (Qiagen Inc., Germantown, MD). Extracted samples were quantified and Illumina Tru-Seq or nano Tru-seq libraries were constructed according to the sample concentration. Samples were sequenced in pools of 24 individuals in 1 lane of an S4 flowcell in an Illumina Novaseq 6000 System with pair-end 150 reads.

### Raw read QC

Reads were aligned with a set of exon sequences to gauge the quality of raw reads before the assembly stage. This step was taken to proactively assess the completeness of the gene benchmarking standard TABASCO^25^ (Test of Absence for Benchmarking Andropogoneae Single-Copy Orthologs). This standard draws from BUSCO^56^ (Benchmarking Universal Single-Copy Orthologs) and encompasses exonic sequences derived from the reference genomes of *Zea mays* subsp. *mays* (B73 RefGen_v5) and *Sorghum bicolor* (v3.1.1). These sequences were curated based on the following criteria:

First, small and large exons (500 bp < x < 4000 bp) were filtered. Then, exons mapping to transposon proteins were removed by aligning sequences against the Viridiplantae (v3.0) transposable element protein domain reference database (REXdb)^57^ using the BLASTx program (v2.9.0)^58^. Next, conserved exons were identified by mapping the previously filtered exon profile against 14 draft Andropogoneae genomes using minimap2 (v2.17)^59^ (parameters *-ax sr*). Exons were removed if they were (1) unaligned and clipped, (2) had a minimal edit distance > 0.1, and (3) were not identified in at least 11 of the 14 reference genomes (∼75%). Finally, the remaining exons were profiled against 58 contig assemblies for conservation using a custom Kotlin script, evaluating how many matches occurred and the degree of CDS conservation.

Exons were removed if less than 70% and greater than 95% were identified in the assemblies (e.g. 0.7 < x < 0.95).

Using this curated profile, raw reads were aligned using minimap2 (v2.17) (parameters *-ax sr*). From these alignments, reads were counted for each exon using similar metrics to those described previously. Raw counts were then adjusted for length correction:

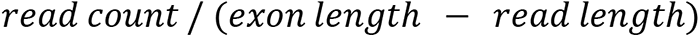

and coverage:

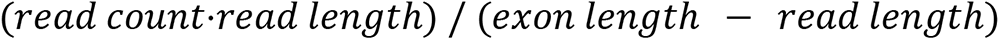

From these metrics, minimum coverage was calculated and used for further assessment.

### Short Read Assembly

Assemblies were generated using Megahit^60^ using a minimum kmer size (--k-min) of 31 and default parameters for the other settings. Other parameters were evaluated through a series of tests on 5 samples previously assembled with ABySS that showed a range of complete genes through TABASCO^25^. We determined that increasing –k-min from 21 to 31 resulted in correctly stitched contigs when compared to other parameters via QUAST^61^ and manual evaluation of genes via MiniMap2^59^. All assemblies were evaluated with TABASCO to evaluate the completeness of a set of close to 6,000 genes in each assembly.

Raw reads were run through the previously described Raw Read QC step, assembled through Megahit, and then evaluated with TABASCO. If an assembly did not pass the QC, or TABASCO step, then that sample was resequenced.

### Gene Model Pre-Processing

For each diverse maize line, we downloaded the GFF and whole-genome FASTA from MaizeGDB. Using a custom Biopython script^62^, we concatenated the 5’ UTR, CDS, and 3’ UTR sequence for each transcript in the GFF, excluding the introns. We then created our own CIGAR-like string of the transcript that allows for easy reconstruction of the gene model. (i.e., GeneID:nL-nC-nT where n denotes the number of basepairs, L leader sequence, C coding sequence, T terminator sequence, and – denoting introns). Annotated transcripts were then aligned to each of the 91 assemblies using minimap2^59^ with parameters --splice and --eqx. This process was parallelized using GNU Parallel^63^. The resulting SAM files were used for generation of the multiple sequence alignments.

### Multiple Sequence Alignment Generation

Each SAM file was parsed using a custom Kotlin script. For each aligned maize transcript, the script counts how many mappings occurred across all species and if there were multiple matches within a species. If multiple matches occurred, the first occurrence was saved. Additionally, within each SAM file, the primary alignments were evaluated for CDS conservation. At least 90% of the read had to be mapped to be saved, otherwise we ignored the read. The script then extracted the alignments of the maize transcript to each Andropogoneae genome and placed them into an individual FASTA file for multiple sequence alignment (MSA) generation. Included in this file was B73 padded with 30 Ns between introns. The output of this script left the first and last 10 basepairs of the introns (partial intron) and padded 10 Ns in the center for each of the other assemblies. In total, over 1.8 million FASTA files were generated for multiple sequence alignment.

The FASTA files were then aligned with MAFFT version 7^64^ with the following parameters:

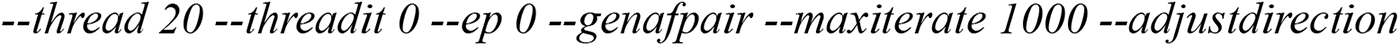

and used in parallel with GNU parallel^63^.

Resulting MSA FASTA files were then processed for downstream inclusion in models. Custom Kotlin scripts were used to split out the leader, coding, and terminator sequences in each MSA. Unique junctions for all transcripts of a single gene were identified and split out into acceptor- and donor-specific MSA files. Additionally, the translation start and stop sites and transcription start and stop sites were extracted and placed into individual MSA files.

### Summary statistics for machine learning models

For each transcript, 4 categories of statistics were calculated based on the MSA to summarize sequence conservation and/or assembly quality of different sequence components (i.e., leader (5’ UTR), coding, terminator (3’ UTR), etc.). The 4 categories include: phyloP conservation score, missingness, reference allele frequency and reference sequence context.

### PhyloP conservation score

We used phast toolkits^65^ to estimate per site phyloP score across 91 *Andropogoneae* species for each transcript when homologous sequences are present and aligned. Briefly, phyloP represents the result of a branch length test against a neutral evolutionary model. Higher absolute scores indicate a significant deviation from neutrality. The direction of deviation is designated by its sign: positive scores denote conservation while negative scores denote acceleration. Considering the variation of neutral substitution rate across the genome, separate neutral models were fitted for each transcript independently based on 4-fold degenerate (4d) sites with the guidance of a species tree constructed from the median pairwise 4d distances across all transcripts. We excluded transcript models with less than 10 informative (polymorphic in at least one pairwise taxa comparison) 4d sites for phyloP calculation due to the lower confidence in the neutral models.

For splicing, translational and transcriptional junction evaluation, phyloP scores of the 20 bp around each unique junction site were included as separate features. For full transcript model evaluation, we used median phyloP score of the coding, leader, terminator and partial intron as input features for model training.

### Missingness

To capture the signatures of presence and absence polymorphism and/or large insertion-deletion polymorphism and/or variation in assembly quality, we calculated the per-site missingness based on the MSA for each transcript model as follows:

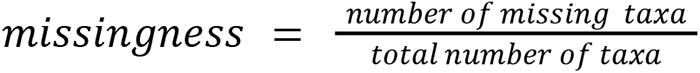

Similarly, for junction evaluation, missingness of the 20 bp around each unique junction site were included as separate features. For full transcript model evaluation, we used median phyloP score of the coding, leader, terminator and partial intron as input features for model training.

### Reference allele frequency

As a separate estimate of sequence conservation, we calculated the frequency of the maize allele among the non-missing taxa (RefFreq) as follows:

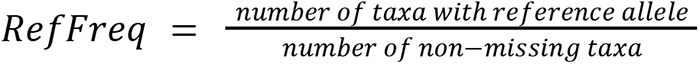

Similarly, for junction evaluation, RefFreq of the 20 bp around each unique junction site were included as separate features. For full transcript model evaluation, we used median phyloP score of the coding, leader, terminator and partial intron as input features for model training.

### Reference sequence context

For splicing and translational junctions, sequence contexts (i.e., splicing motifs and translation start/stop codons) are crucial. We incorporated sequence contexts into our junction evaluation by numerically encoding each nucleotide of the 20 bp junction reference sequence (A:1, T:2, C:3, G:4) as features for model training.

### XGboost model for splicing and translation junction evaluation

Junction MSAs of 20 bp sequence around all unique junction sites (10bp upstream and 10bp downstream the splicing donor, acceptor, translation start and stop sites) were extracted from the intron-shortened transcript MSAs. We considered the junctions in the core gene sets across maize genomes^22^ as positive (valid junctions). To generate negative (mal-functional) junction sequences, a balanced subset of junctions from non-core genes were subjected to a customized mutator function with a mutation frequency X, respectively, where X ∼ Pois (lambda=1) (FigS9). Briefly, the mutator function randomly swaps N pairwise positions in a junction MSA to break motif signatures while maintaining the overall phylogenetic relationship between taxa (FigS9).

Summary statistics were calculated per site for all junction MSAs (4 categories x 20 sites; 80 features in total) and used as the features to learn the quality of an evolutionary functional junction. Given the difference in splicing motifs, acceptors and donors were processed independently for independent model construction.

Four XGboost models^28^ were trained for junction evaluation for splicing donor, acceptor, translation start and stop sites separately. In both cases, we did a 1-to-1 split for the training and validation datasets (N = ∼190,000 for splicing junction models and ∼35,000 for translation junction models). The XGboost models were trained on a total of 80 features with a maximum tree depth of 6, learning rate (eta) of 0.1 and 1000 runs of iteration in the R environment.

Precision-Recall (PR) and Receiver operating characteristic (ROC) curves were plotted. The area under curve (AUC) was used to evaluate the models.

### XG-Boost model for transcription Start/Stop junction evaluation

MSAs for the 20bps around the annotated transcription start and stop sites were created. For the transcription start site, we considered transcripts that had CAGE^26^ support to be positive, and those without support to be negative. We additionally followed a similar shuffling procedure as outlined for the splicing/translation Junction models to simulate negatives. Our positives for the transcription stop site were transcripts that had PolyA support^27^, and our negatives ones without support or again simulated shuffling of the stop site. Similarly, the XGboost models were trained on a total of 80 features with a maximum tree depth of 6, learning rate (eta) of 0.1 and 500 runs of iteration in the R environment. The models were evaluated on a set of 5,000 withheld genes. The same shuffling process was included during the evaluation. Precision-Recall (PR) and Receiver operating characteristic (ROC) curves were plotted. The area under curve (AUC) was used to evaluate the models.

### LSTM for mRNA model

We generated an input matrix for each transcript that consisted of summary statistics for each exon and intron, as described above. Additionally, we included the XG-Boost model scores for the translation start site, donor sites, acceptor sites, and translation stop site in the respective column for the input matrix. Each column represented an exon or an intron, resulting in variable lengths depending on the transcript, with a fixed number of rows. The input matrices were then converted into a ragged tensor, batched into sets of 64, and fed into a LSTM model using Keras^66^. The LSTM model consisted of a bidirectional LSTM layer followed by three dense layers (the final containing a sigmoid activation function) to get a single output ranging between zero and 1. A leave one chromosome out (LOCO) training strategy was used, where all transcripts across NAM were grouped together by chromosome, training on nine and leaving out one. The training set consisted of all core B73 transcripts and all noncore B73 transcripts, which were supplemented with additional NAM noncore transcripts that were selected at random in order to fully balance the training set. 80% of the total number of core and noncore transcripts were used. The validation set was kept proportional to the actual frequency of core/noncore for that chromosome, and equaled 20% of the remaining transcripts. The test/prediction set was all NAM transcripts for the left-out chromosome.

### XG-Boost for proteins

ProtTrans^20^ is a transformer-based protein language model which was trained with 2.123 billion protein sequences from Bacteria, Viruses, Archaea and Eukaryota. Using a self-supervised approach, it treats protein sequences as sentences and amino acids as tokens, aiming to predict masked amino acids. The model captures protein folding patterns and evolutionary conservation via 1,024-dimensional embeddings. To fully leverage the protein structure and conservation learned by ProtTrans, we trained an XG-Boost model with 1,024-dimensional embeddings. In addition to using dispensable and private genes as negatives, we also generated simulated intron retention and exon exclusion genes as negatives to learn the differences between core and non-core proteins as much as possible. We used the same LOCO training strategy as the mRNA model, the final predictions for each transcript are a combination of 10 LOCO models.

### reelGene model

Scores from the LSTM (mRNA model) and protein XG-Boost (protein model) were averaged within each transcript. UTR XG-Boost models (start model and stop model) were also calculated, but kept separate from the main reelGene model.

### Scripts and Data Availability

All data and scripts are available upon request, and will be deposited in our repository upon publication.

## Acknowledgments

This work is funded by the USDA-ARS and NSF PanAnd Grant Award #1822330. A.J.S. is supported by NSF GRFP DGE – 2139899. T.H.F. is supported by USDA NIFA AFRI predoctoral fellowship (TF: 2022-67011-36564). We thank Jose A Valdes Franco for his help with minimap2 parameter tuning, Kay McNeary for designing the reelGene logo, George Stack for coming up with the reelGene acronym, Qi Sun for helping with the shortread assembly pipeline, and Subham Sahoo for machine learning consulting.

