## Supplemental Figures for "Fishing for a reelGene: evaluating gene models with evolution and machine learning"

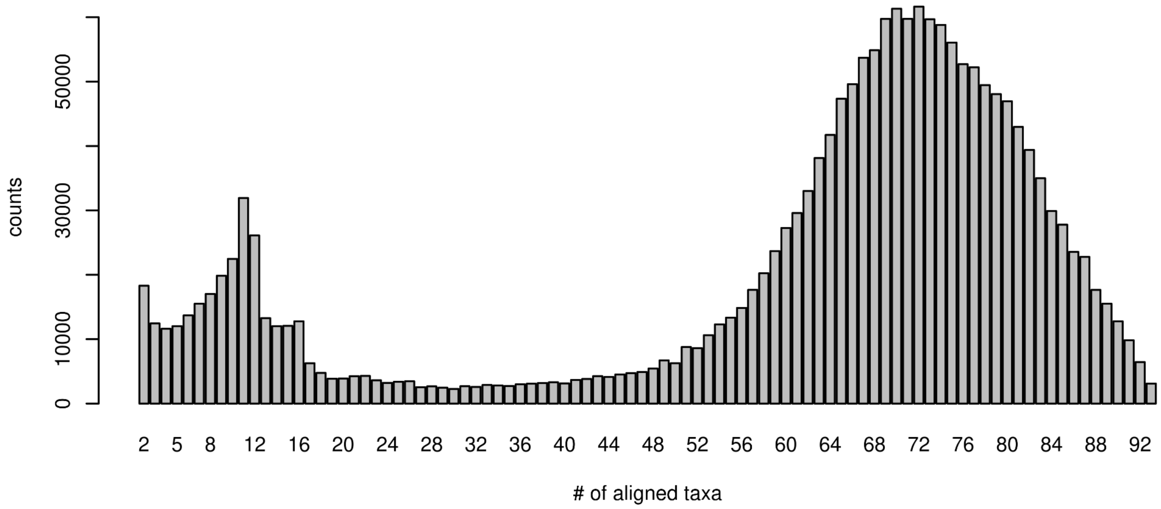


**Figure S1. Numbers of aligned taxa for each maize gene model.** Among all 1.8 million gene models annotated in 26 different maize lines, we found alignable homologous sequences in 62 taxa on average, with a mode at 72. In the second mode at 10, most of the gene models are *Zea*-specific.

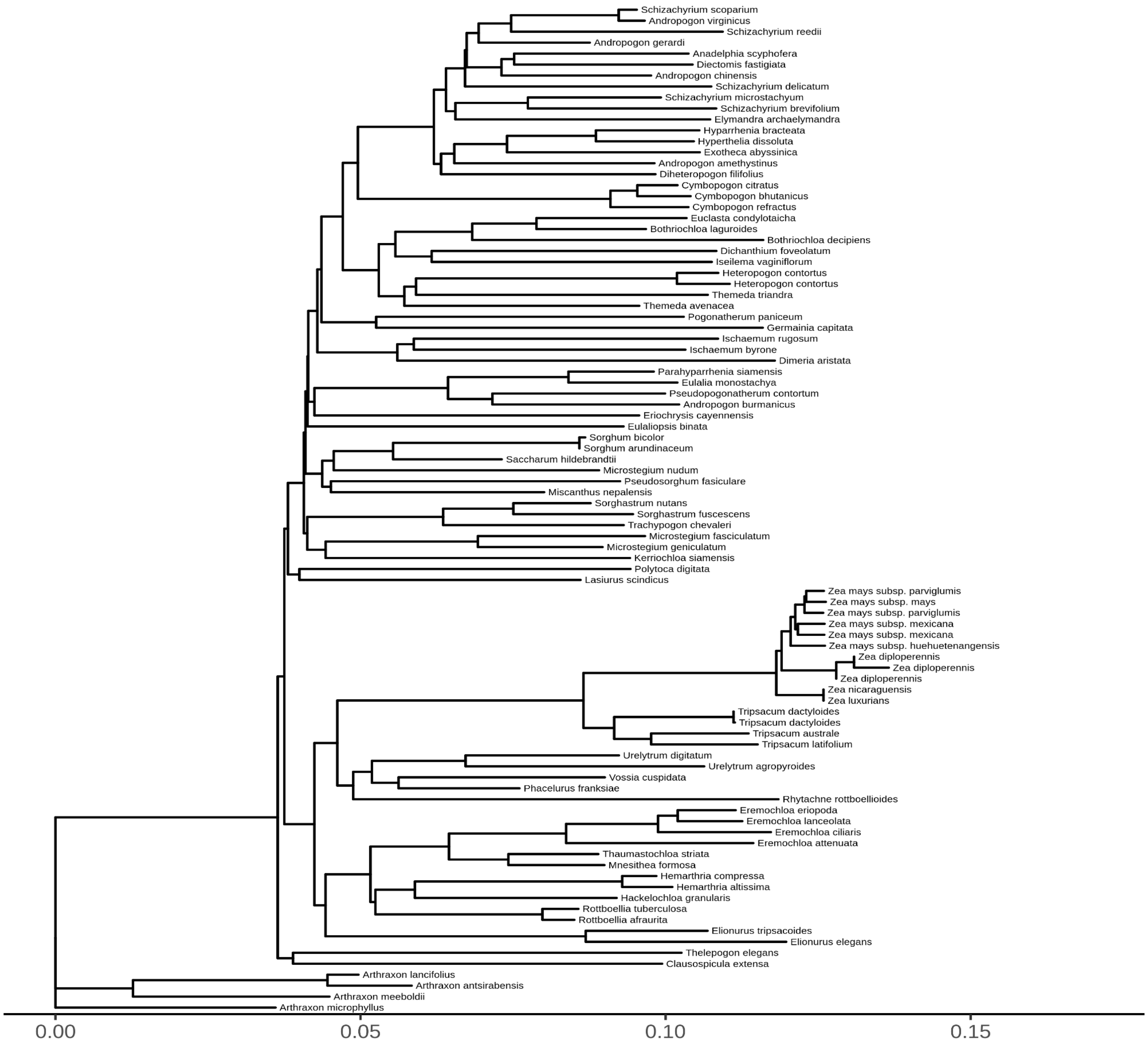


**Figure S2. Species phylogeny of the 92 Andropogoneae taxa.** We calculated genetic distance matrices based on the 4-fold degenerate sites for each gene annotated in the maize reference genome (B73v5). Neighbor-joining phylogeny was constructed based on the median distance across all genes. The tree is rooted on *Arthraxon microphyllus*.

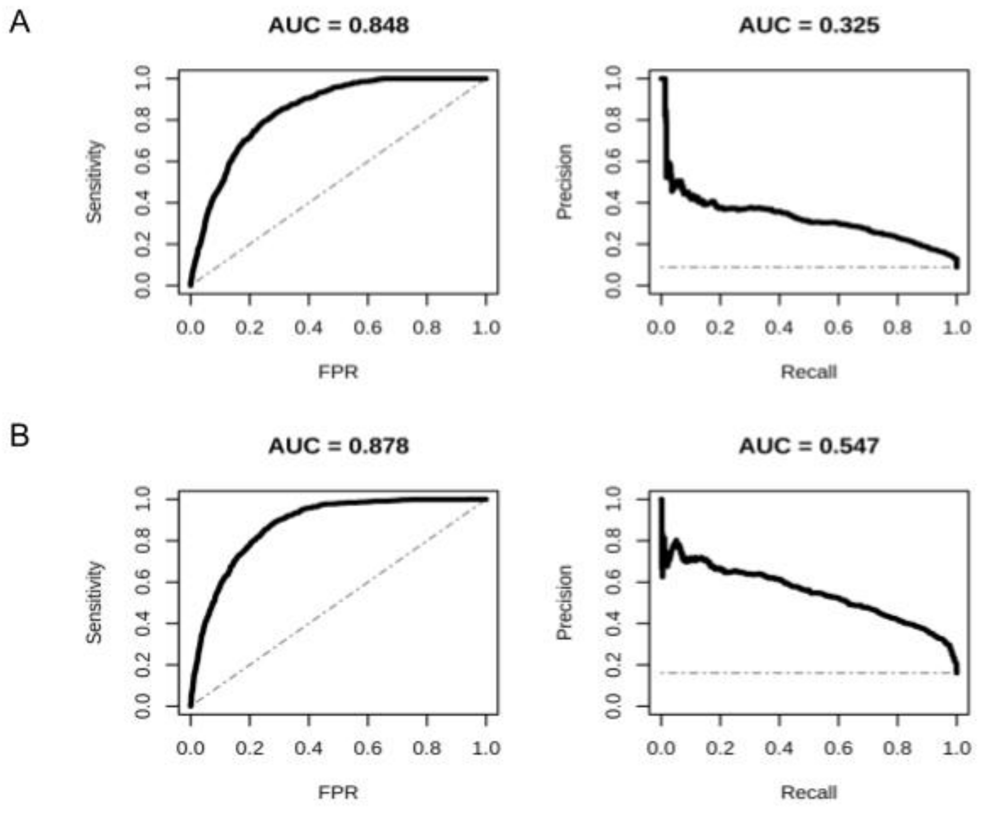


**Figure S3. Transcript boundary models performance.** The model performance for the transcriptional start (A) and stop (B) junctions. The evaluation of the models were done on 5,000 withheld genes with positional swapping in the negatives.

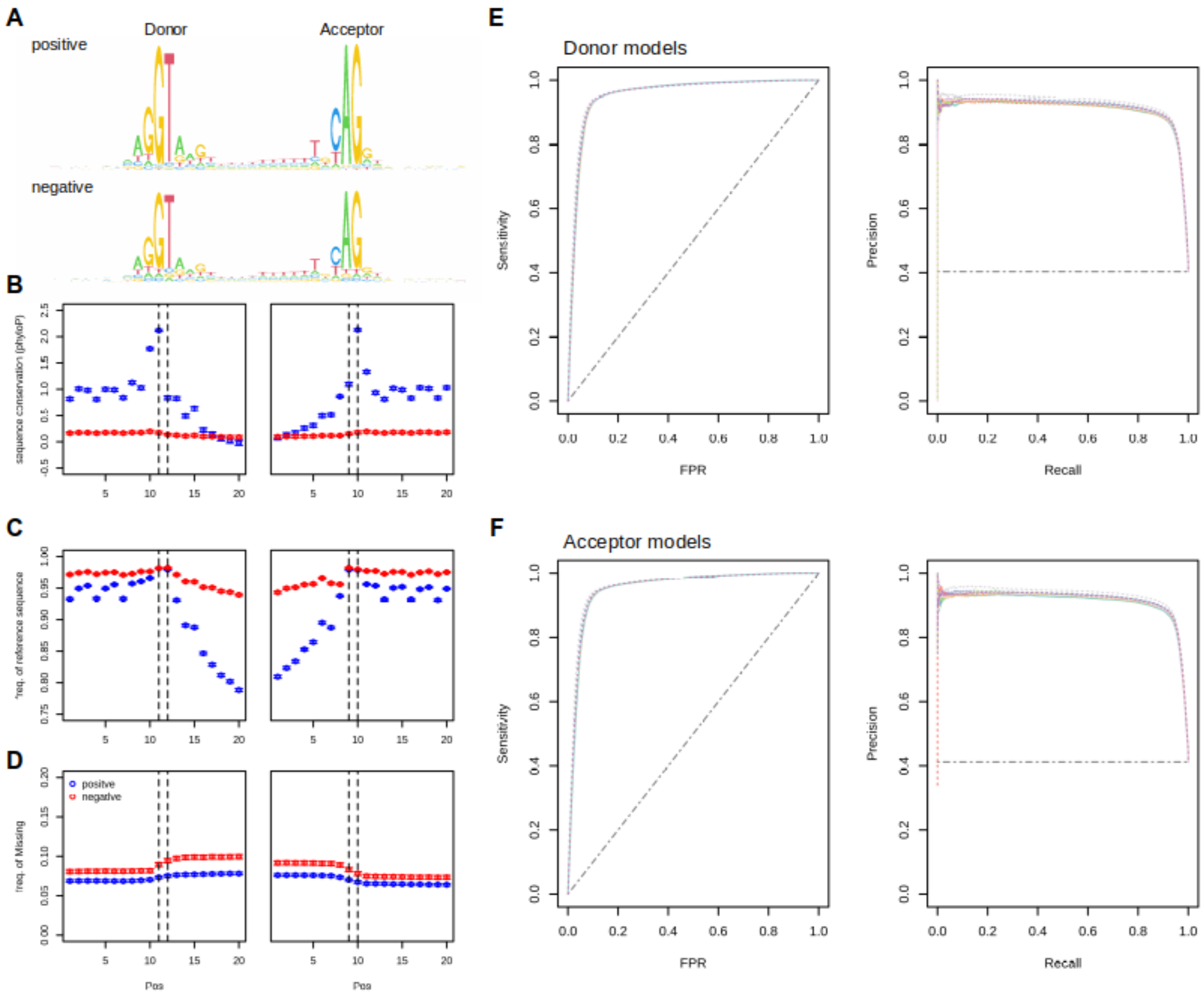


**Figure S4. XGboost models for splicing junctions. A-D**. The distribution of different per-site summary statistics around the positive splicing junctions (junctions in core genes) and the negative ones (junctions in dispensable and private genes with positional swapping). These statistics were the input of the XGboost models. **E & F**. The performance of the models across 10 leave-one-chromosome-out (LOCO) training-validation cycles (denoted by different colors).

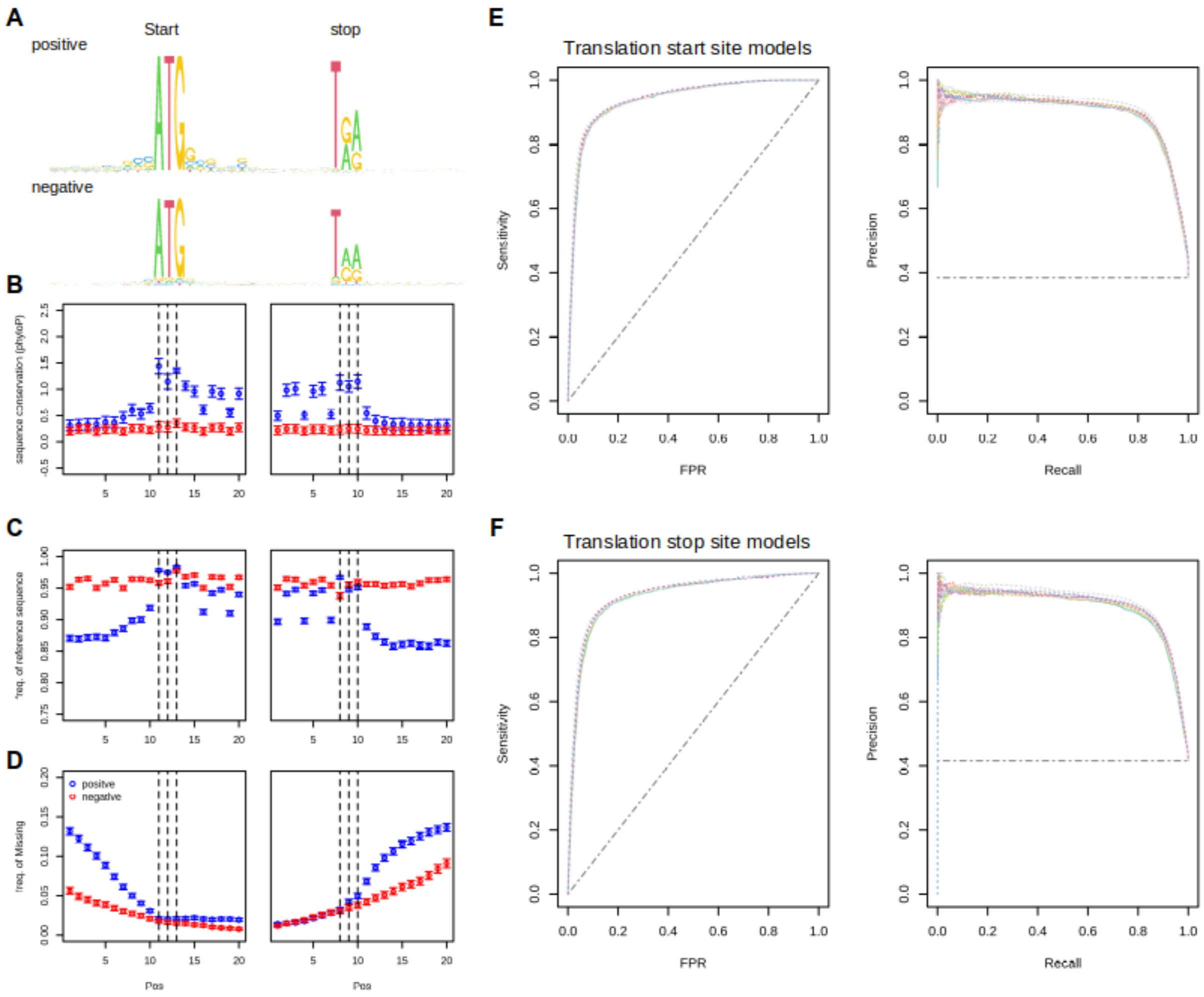


**Figure S5. XGboost models for translational junctions. A-D**. The distribution of different per-site summary statistics around the positive translational junctions (junctions in core genes) and the negative ones (junctions in dispensable and private genes with positional swapping). These statistics were the input of the XGboost models. **E & F**. The performance of the models across 10 leave-one-chromosome-out (LOCO) training-validation cycles (denoted by different colors).

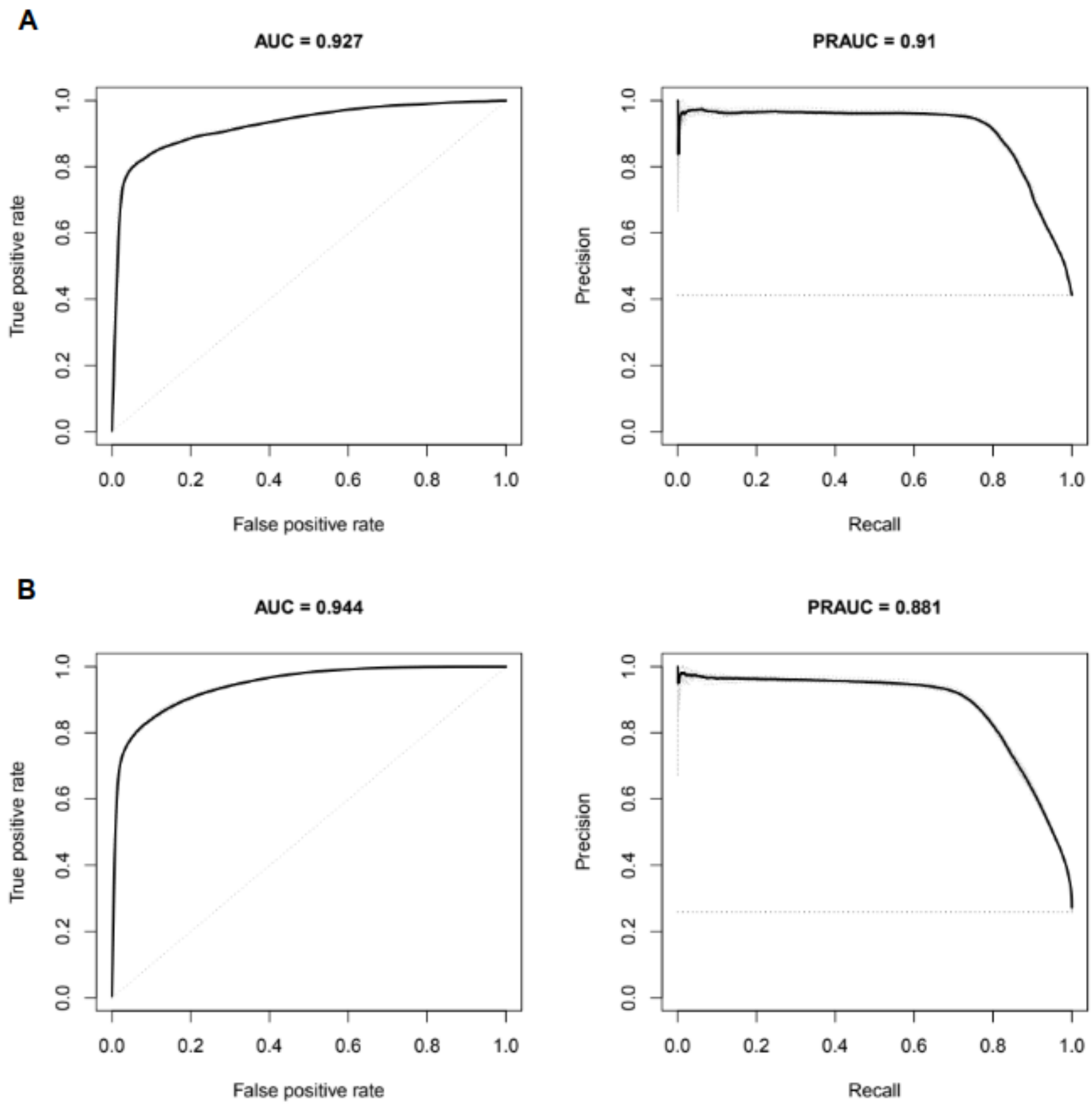


**Figure S6. Model performance of the mRNA (A) and protein (B) models.** The performance of the models across 10 leave-one-chromosome-out (LOCO) training-validation cycles (gray dash lines) were shown with the average as a black solid line.

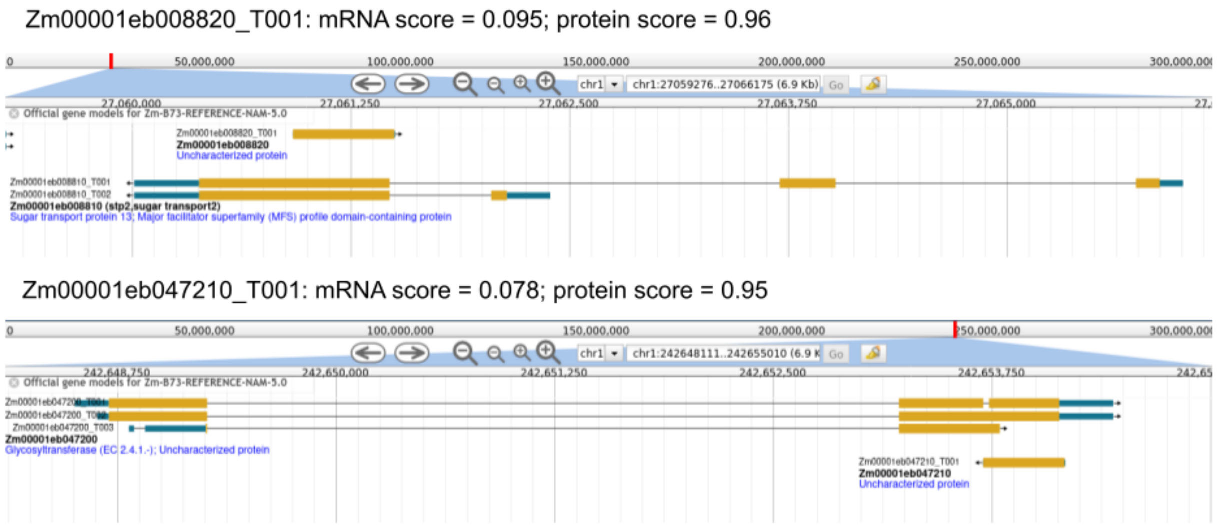


**Figure S7. Examples of mis-annotated antisense RNA with high protein score but low mRNA score.** We provide two examples of annotated gene models with high protein scores but low mRNA scores where they seem more likely to be antisense non-coding RNAs but mis-annotated as protein-coding genes. Our mRNA model is able to detect such mis-annotation as it learns and checks the sequence context for all essential junctions of a protein-coding gene.

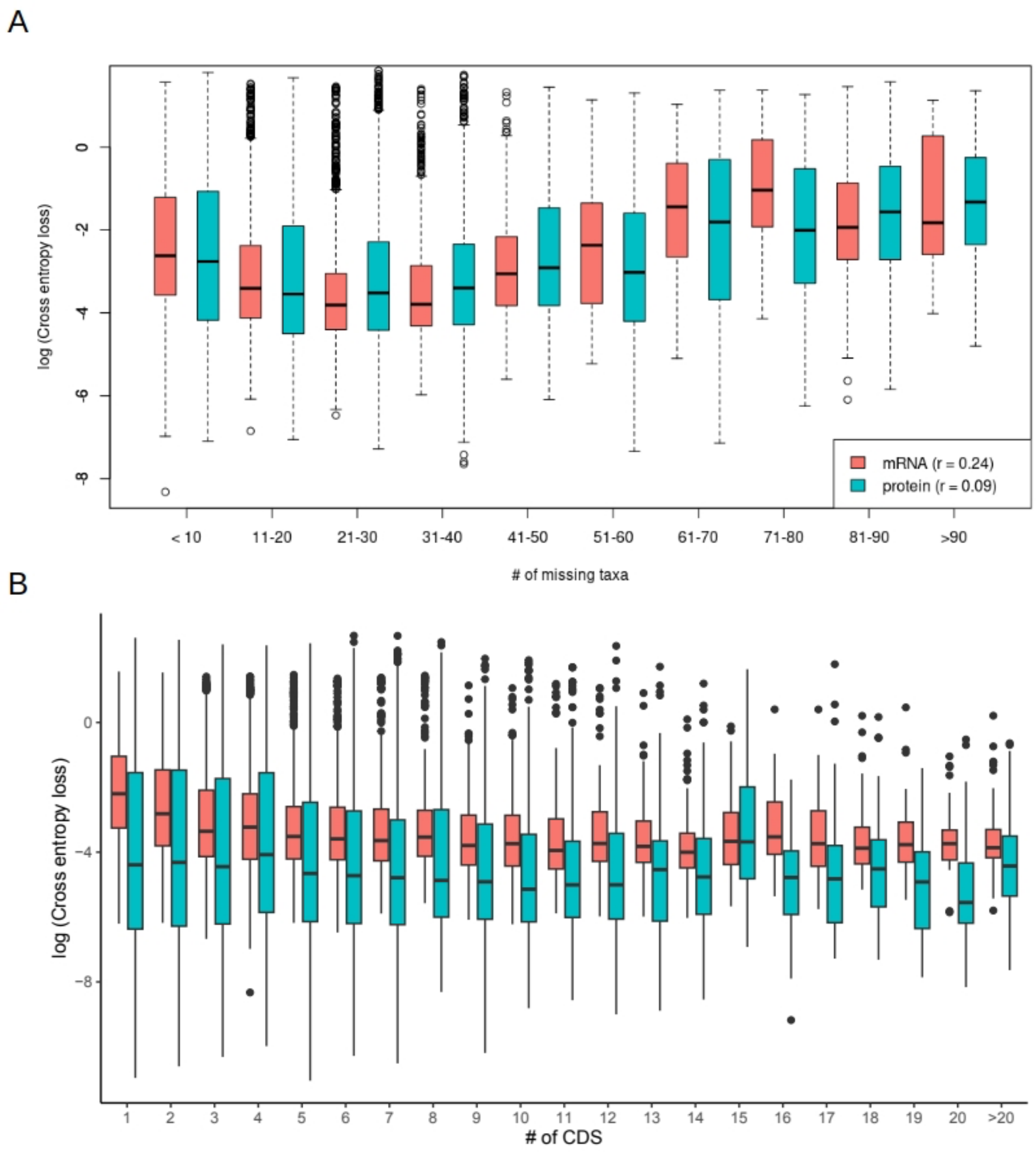


**Figure S8. The relationship of classification error (higher cross entropy loss) and the completeness of alignment (A) and the numbers of CDS exons (B) within the mRNA and protein models.** We evaluated the cross-entropy loss of the mRNA and protein models in relation to the number of Andropogoneae taxa missing from each transcript multiple sequence alignment (A) and the number of CDS exons (B) in each transcript. As expected, mRNA showed higher reliance on the completeness of alignment and made more mistakes on gene models with fewer CDS exons.

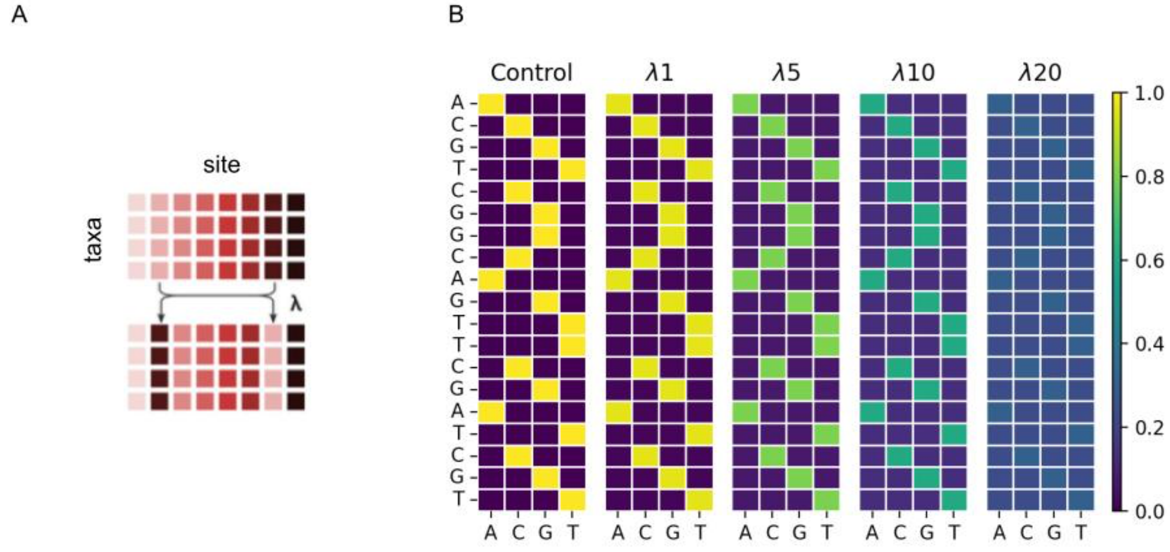


**Figure S9. Illustration of the positional swapping process to introduce false junctions.** To add more true negative signals for junction model training, we included a swapping process to break down the positional conservation patterns around key junctions (i.e., translational start and stop sites and splicing donor and acceptor sites) of a transcript. **A**. Pairwise swapping of the sites occurred as a Poisson process, where sites were swapped, but the relationship between taxa remained unchanged. **B**. With increasing lambda of the Poisson distribution, the patterns of site conservation declined. In the formal analysis, lambda of 1 was used.
